# Strain-level identification of bacterial tomato pathogens directly from metagenomic sequences

**DOI:** 10.1101/777706

**Authors:** Marco E. Mechan Llontop, Parul Sharma, Marcela Aguilera Flores, Shu Yang, Jill Pollock, Long Tian, Chenjie Huang, Steve Rideout, Lenwood S. Heath, Song Li, Boris A. Vinatzer

**Author notes:** These authors contributed equally (Marco E. Mechan Llontop, Parul Sharma, Marcela Aguilera Flores). Corresponding authors: Boris A. Vinatzer and Song Li, E-mail addresses.

## Abstract

Routine strain-level identification of plant pathogens directly from symptomatic tissue could significantly improve plant disease control and prevention. Here we tested the Oxford Nanopore Technologies (ONT) MinION™ sequencer for metagenomic sequencing of tomato plants either artificially inoculated with a known strain of the bacterial speck pathogen *Pseudomonas syringae* pv. *tomato* (*Pto*), or collected in the field and showing bacterial spot symptoms caused by either one of four *Xanthomonas* species. After species-level identification using ONT’s WIMP software and the third party tools Sourmash and MetaMaps, we used Sourmash and MetaMaps with a custom database of representative genomes of bacterial tomato pathogens to attempt strain-level identification. In parallel, each metagenome was assembled and the longest contigs were used as query with the genome-based microbial identification Web service LINbase. Both the read-based and assembly-based approaches correctly identified *Pto* strain T1 in the artificially inoculated samples. The pathogen strain in most field samples was identified as a member of *Xanthomonas perforans* group 2. This result was confirmed by whole genome sequencing of colonies isolated from one of the samples. Although in our case, metagenome-based pathogen identification at the strain-level was achieved, caution still needs to be exerted when interpreting strain-level results because of the challenges inherent to assigning reads to specific strains and the error rate of nanopore sequencing.

## Introduction

Early detection of plant disease outbreaks and accurate plant disease diagnosis are prerequisites of efficient plant disease control and prevention (Tinivella et al. 2008). In many cases, an experienced plant pathologist can quickly diagnose a disease based on symptoms. However, visual diagnosis does not identify the causative agent at the strain-level. For example, three different strains of the plant pathogen *Pseudomonas syringae* pathovar (pv.) *tomato* (*Pto*) cause indistinguishable bacterial speck disease symptoms in tomato (Cai et al. 2011). Sometimes, visual diagnosis cannot even identify a pathogen at the species level. For example, four different species of the genus *Xanthomonas* cause indistinguishable bacterial spot disease symptoms on tomato (*Solanum lycopersicum*) leaves (Jones et al. 2004). Note that in this article, we use the term “strain” as an intraspecific, monophyletic group of bacteria, which have a very recent common ancestor and are thus genotypically and phenotypically more similar to each other than to other members of the same species (Dijkshoorn et al. 2000). To avoid confusion, we use the term “isolate” instead of “strain” when referring to a pure culture of bacteria isolated on a specified date at a specified geographic location from a specific plant.

While most disease control measures may be the same for different pathogen strains or species, depending on the precise identity of the pathogen, additional control measures may need to be undertaken. For example, different strains of the same pathogen species may have different host ranges. Therefore, it may be necessary to avoid certain crop rotations or to eliminate certain weeds depending on the identity of the strain that causes a disease and its specific host range. In the case of *Pto*, strain T1 causes disease only in tomato while strain DC3000 causes disease in tomato and in leafy greens of the family *Brassicaceae* (Yan et al. 2008). Strain DC3000 could thus spread from tomato fields to leafy green fields, cause disease in a leafy green planted after tomato, and/or survive in weeds that belong to the Brassicaceae family. In other cases, identifying a pathogen to strain level could even trigger eradication procedures to stop further spread of the disease. For example, this would happen if the select agent *Ralstonia solanacearum* Race 3 Biovar 2 were to be identified as the causative agent of bacterial wilt disease outbreak in the USA (Williamson et al. 2002). Fast strain-level plant pathogen identification would thus add significant value to plant disease diagnostics.

Many molecular tools have been developed over the years for pathogen identification. They all have their strengths and weaknesses (Fang and Ramasamy 2015). Many of them depend on a pure pathogen culture and thus require lengthy procedures to isolate and culture the pathogen from the plant tissue. Moreover, many of them cannot identify pathogens at the strain level. Gene sequence-based techniques, such as multilocus sequence typing/analysis (MLST/A) (Almeida et al. 2010), can identify a pathogen to strain-level but usually require pure cultures. Moreover, gene sequence-based techniques depend on previous species-level identification because different species require different primers to amplify the genes to be sequenced by polymerase chain reaction (PCR), for example see (Rees-George et al. 2010). One alternative gene-based method is to amplify the 16S rRNA gene directly from DNA extracted from plant tissue and to identity the putative pathogen based on its 16S rRNA sequence. We have recently tested this method but not found it to be suitable because of its low resolution (Mechan-Llontop et al. 2019).

Whole genome sequencing (WGS) does not require PCR and strain-level identification is now routine practice in the surveillance of food-borne pathogen outbreaks in several countries (Nadon et al. 2017). With the drop in sequencing cost and development of genome databases that contain strain-level classification of plant pathogens, WGS now represents a real possibility in plant disease diagnostics. For example, LINbase at linbase.org (Tian et al. 2019) contains precise genome-based circumscriptions for many bacterial plant pathogens from the genus level to the strain level. Genome sequences of unknown isolates can be identified as members of circumscribed plant pathogens based on how similar they are at the whole genome level, measured as Average Nucleotide Identity (ANI) (Konstantinidis and Tiedje 2005), to the other members of these taxa. However, the limitation of WGS is its dependence on pure cultures.

Metagenomic sequencing consists in extracting DNA directly from plant tissue followed by sequencing all DNA present in the sample. Compared to WGS, the two main advantages of this approach are that (1) it is much faster because it does not require lengthy pathogen isolation and culturing procedures; and (2) it does not require much prior knowledge about the pathogen since any pathogen, besides RNA viruses, can be detected with this method. However, the main challenge of this approach is that the obtained DNA sequences also contain host plant sequences and microbe sequences that do not belong to the pathogen. Therefore, obtaining sufficient sequences of the causative agent and identifying the causative agent among all the other potential causative agents present in the same plant requires optimized experimental methods for DNA extraction and sequencing and optimized algorithms and genome databases for precise pathogen identification.

The sequencing method that is currently most attractive for metagenomics-based pathogen identification is nanopore sequencing with the Oxford Nanopore Technologies (ONT) MinION™ device (Jain et al. 2016). The main strengths of this method are that (1) DNA can be prepared for sequencing with relatively short protocols (from a few hours to less than an hour; https://community.nanoporetech.com), (2) the MinION™ sequencer is not much larger than a USB stick and can be used with a desktop or a laptop computer in the lab or even in the field, it provides the first sequencing results within minutes from the start of a sequencing run, and the output can reach over 10 gigabases of DNA sequences (more than 1000 times the size of an individual bacterial genomes) after 48 hours (MinION brochure 2019a). However, the major weaknesses are (1) the high sequencing error rate of approximately 10% (Tedersoo et al. 2019; Loit et al. 2019) and (2) that the sequencing hardware only works once at full capacity limiting reuse (MinION brochure 2019b).

Metagenomic sequencing with the MinION™ has already been used on several crops for identification of various pathogens (Chalupowicz et al. 2019) using ONT’s software WIMP (Juul et al. 2015) and on wheat to identify various fungal pathogens (Hu et al. 2019) using the sequence alignment tool BLASTN (Camacho et al. 2009) in combination with custom databases. The MinION™ has also been used for plant pathogen detection and identification starting from extracted RNA or DNA in combination with general or specific primers to increase the quantity of input for the MinION™ (Loit et al. 2019; Badial et al. 2018). However, in none of these studies, was strain-level identification attempted directly from sequencing metagenomic DNA without prior amplification.

Here we tested the MinION™ with tomato plants artificially inoculated with different strains of *Pseudomonas syringae*, including isolates of the *Pto* strains T1 and DC3000 (Cai et al. 2011), and with plants from tomato fields showing symptoms of natural infection with bacterial spot for which we did not know the *Xanthomonas* species that caused the infection. We then explored the precision of identification that can be achieved when using ONT’s WIMP software, Sourmash (Brown and Irber 2016), and MetaMaps (Dilthey et al. 2019) in combination with default and custom reference databases. We also assembled metagenomic sequences into contigs and identified contigs in combination with BLASTN (Camacho et al. 2009) and in combination with the LINbase Web service for genome-based microbial identification (Tian et al. 2019).

## Materials and Methods

### Laboratory-infected tomato plants

Seeds of tomato (*Solanum lycopersicum*) ‘Rio Grande’ were germinated in potting mix soil (Miracle-grow, OH, USA) under laboratory conditions with a long day period (16-h photoperiod) and infected at 4 weeks of age. *Pto* isolate K40 (belonging to strain T1), *Pto* isolate DC3000 (belonging to strain DC3000) (Cai et al. 2011), *P. syringae pv. syringae* B728a (Feil et al. 2005), and *P. syringae* 642 (Clarke et al. 2010) were grown in King’s B solid medium at 28°C for 24 hours. Isolate *Pto* K40 was suspended at a concentration corresponding to an OD600 of 0.001 in 10 mM MgSO4 for single-strain inoculation. For the mixed-strain inoculation, all four isolates were suspended at an OD600 of 0.001 in 10 mM MgSO4 and pooled together in equal amounts before inoculation. Silwet L-77 was added to bacterial suspensions (0.025% vol/vol) to facilitate bacterial infection. Plants were placed in ziplock plastic bags for high humidity conditions for 24 hours before inoculation. After plants were spray-inoculated with 10 ml of bacterial suspensions, they were placed back into the plastic bags for another 24 hours. Plants were processed for DNA extraction three days later. Inoculation with 10mM MgSO4 was included as a mock treatment.

### Naturally infected tomato plants

Five tomato plants with bacterial spot symptoms, one plant with symptoms of Septoria leaf spot, and one plant without symptoms were collected on August 10, 2018, on the Eastern Shore of Virginia (Accomack and Northampton counties) and shipped overnight to the Virginia Tech campus in Blacksburg, VA, where they were processed for DNA extraction. Another set of plants with bacterial spot symptoms were collected in May, 2019. Bacteria were isolated from symptomatic leaves on King’s medium B. Plants and plates were shipped to the Virginia Tech campus overnight where plants and bacterial colonies were processed for DNA extraction.

### DNA extraction

All plant samples used for DNA extraction are listed in Table 1. DNA extraction was performed according to (Ottesen et al. 2013) with the following modifications. Briefly, wearing gloves, the top of each plant sample (6 to 10 leaves from the top with or without stems) was collected using clippers. The weight of samples was between 5 to 10 grams. After removing all the dirt from the plant surface by shaking vigorously, each sample was placed in a 6-1/2”× 5-7/8” Ziploc® bag together with 300 ml sterilized double-distilled water (DDW). Samples were sonicated for 15 minutes using a Branson 1510 Ultrasonic Cleaner. DNA was extracted with DNeasy® PowerWater® Kit (QIAGEN; Catalog # 14900-50-NF). All steps for DNA extraction were performed according to the kit’s specifications, except that after adding 1 mL of the kit’s solution PW1, the tube was incubated at 65°C for 15 minutes and then vortexed for 20 minutes.

**Table 1.** Description of samples used in this study.

| Sample Name | Short description | DNA concentration of samples (ng/ul) | Fraction of flow cell used | # reads base-called | Total length of reads base-called | % of reads classified as bacteria (based on WIMP) | Mean read length in bp | Max read length in bp | % reads >1000bp |
| --- | --- | --- | --- | --- | --- | --- | --- | --- | --- |
| L-K40 | Tomato inoculated with *Pto* K40 in the laboratory | 325.2 | 1 | 1,377,617 | 4.18 Gb | 89% | 3,037 | 66,015 | 64% |
| L-mix | Tomato inoculated with four *P. syringae* strains in the laboratory | 450.4 | 1 | 1,006,978 | 4.16 Gb | 95% | 4,130 | 67,174 | 74% |
| L-mock | Non-inoculated tomato plant grown in the laboratory | 33.6 | 1/7 | 82,412 | 103.22 Mb | 8% | 1,252 | 19,754 | 40% |
| L-culture-mix | Equal mix of 4 *P. syringae* strains grown in liquid culture | 147.5 | 1/6 | 54,124 | 155.93 Mb | 93% | 2,880 | 76,060 | 39% |
| F1-bs | Tomato field sample with symptoms of bacterial spot | 562 | 1/7 | 137,497 | 588.50 Mb | 81% | 4,280 | 55,436 | 73% |
| F2-bs | Tomato field sample with symptoms of bacterial spot | 500.2 | 1/7 | 90,185 | 498.68 Mb | 80% | 5,529 | 65,598 | 74% |
| F3-bs | Tomato field sample with symptoms of bacterial spot | 332.5 | 1/7 | 100,956 | 423.16 Mb | 78% | 4,191 | 59,405 | 68% |
| F4-bs | Tomato field sample with symptoms of bacterial spot | 319.8 | 1/7 | 74,615 | 289.36 Mb | 81% | 3,878 | 51,268 | 70% |
| F5-Septoria | Tomato field sample<br>with symptoms of<br>Septoria leaf spot | 75.8 | 1/7 | 73,432 | 226.721 Mb | 50% | 3,087 | 43,967 | 59% |
| F6-healthy | Tomato field sample<br>with no symptoms | 29.1 | 1/7 | 35,923 | 66,58 Mb | 31% | 1,853 | 29,617 | 46% |
| F7-bs | Tomato field sample<br>with symptoms of<br>bacterial spot | 331.8 | 1/7 | 118,391 | 432.08 Mb | 75% | 3,649 | 48,335 | 64% |
| F8-bs | Tomato field sample<br>with symptoms of<br>bacterial spot | 154.2 | 1/2 | 106,059 | 371.84 Mb | 70% | 3,505 | 33,472 | 71% |

DNA from isolated bacteria was extracted with the Gentra® Puregene® Cell and Tissue Kit (Gentra Systems; Catalog # D5000). All steps for DNA extraction were performed according to the Gram-negative Bacteria protocol, except that cells were collected in 1 mL of sterilized DDW in a 1.5 ml microcentrifuge tube for the lysis step. For both extraction procedures, the concentration and purity of DNA was measured using a Thermo Scientific™ NanoDrop™ One^C^ Spectrophotometer.

### DNA library preparation

Library preparation was performed according to the ‘1D Native barcoding genomic DNA protocols (EXP-NBD104, EXP-NBD114, and SQK-LSK108 or SQK-LSK109) provided by ONT. Sequencing libraries were prepared using the Ligation Sequencing Kit (ONT Ltd.; SQK-LSK109). For each run, NEBNext® Ultra™ II End Repair/dA-Tailing Module (New England Biolabs, Inc.; Catalog # E7546S) was used for DNA repair and end-prep for each sample. Repaired DNA was cleaned up by 1.5 volumes of AMPure XP beads, washed on a magnetic rack using freshly made 70% ethanol, and eluted with 25 μL nuclease-free water. 22.5 μL elute was used for barcoding by mixing with the Blunt/TA Ligase Master Mix (New England Biolabs, Inc.; Catalog # M0367S) and Native Barcode (Oxford Nanopore Technologies Ltd.; Native Barcoding Expansion Kit EXP-NBD104), followed by another wash step using 1.5 volumes of AMPure XP beads, and DNA was eluted in 26 μL nuclease-free water. Equimolar amounts of barcoded DNA were then pooled into a 1.5 mL microcentrifuge for ligation. Adapter ligation was performed by mixing the pooled barcoded sample with Adapter Mix (Oxford Nanopore Technologies Ltd.; SQK-LSK109), NEBNext® Quick Ligation Reaction Buffer (New England Biolabs, Inc.; Catalog # B6058S) and Quick T4 DNA Ligase (New England Biolabs, Inc.; Catalog # M2200S). Ligated DNA was cleaned up by one volume of AMPure XP beads, washed on a magnetic rack using Long Fragment Buffer (Oxford Nanopore Technologies Ltd.; SQK-LSK109), and eluted with 15 μL Elution Buffer (Oxford Nanopore Technologies Ltd.; SQK-LSK109).

Sequencing reactions were performed independently for each run on a ONT MinION™ flow cell (FLO-MIN106 R9 Version) connected to a Mk1B device (ONT Ltd.; MIN-101B) operated by the MinKNOW software (latest version available). Each flow cell was primed with the priming buffer prepared by mixing 30 μL Flush Tether (ONT Ltd.; EXP-FLP001) with a tube of Flush Buffer (ONT Ltd.; EXP-FLP001). 12 μL of the final library mixed with Sequencing Buffer (ONT Ltd.; SQK-LSK109) and Library Loading Beads (ONT Ltd.; SQK-LSK109) was loaded onto the SpotON sample port of the flow cell in a dropwise fashion. The sequencing run was stopped after 48 hours.

### Illumina genome sequencing and assembly

Genomic DNA from isolated bacteria was used to prepare 350bp insert DNA libraries and sequence on an Illumina platform PE150 at Novogene Corporation Inc (Sacramento, CA). FastQC was used to assess the quality of the raw sequencing data (Andrews 2010). Adapter-trimming was performed using BBduk with the parameters ‘k=23, mink=9, hdist=1, ktrim=r, minlength=100’ (Bushnell 2015). Unicycler v0.4.7 with default parameters was used to *de novo* assemble the bacterial genomes (Wick et al. 2017).

### Read-based metagenomic analysis

#### Guppy

For all samples, the Fast5 files containing raw reads were base-called with the base-calling ONT software Guppy (v3.3.2), which uses neural networks to translate raw signals into DNA sequences in fastq format (available via https://community.nanoporetech.com).

#### What’s in my pot? (WIMP)

The ONT workflow WIMP (v2019.7.9), which uses Centrifuge (Kim et al. 2016) to assign taxonomy to reads in real-time, was used for species level identification in all samples.

#### Sourmash

Sourmash, a command-line tool used for k-mer based taxonomic classification for genomes and metagenomes, computes MinHash sketches to create signatures of DNA sequences which are then used to assign taxonomic annotations. The *gather* function in this software was used for taxonomic classification at the species- and strain-level. For species-level classification, the default Genbank LCA database (v.2018.03.29, k=31) containing 100,000 microbial genomes was used. For strain level-classification, a custom library with 245 microbial genomes representative of tomato plant pathogens and close relatives was used. A complete list of genomes used in the custom reference library is provided in Supplementary Table 1. For all samples, signatures were computed at 31 k-mer size (for species level) and 51 k-mer size (for strain level) and abundance filtering was performed to exclude k-mers with an abundance of 1 (Brown and Irber 2016). Sourmash was run on Virginia Tech’s High Performance Computing system, Advanced Research Computing (ARC), with 32 cores and 128GB memory.

#### MetaMaps

Metamaps (Dilthey et al. 2019) was used for taxonomic classification at the species-level using the miniSeq+H database, which includes more than 12,000 microbial genomes and is included with the software package. For strain-level classification, the custom library described above for Sourmash was used. However, the list of genomes was reduced to 149 to include only those genomes that had NCBI taxonomy IDs as per a prerequisite for Metamaps. MetaMaps was also run on Virginia Tech’s High Performance Computing system, Advanced Research Computing (ARC), with 32 cores and 128GB memory.

#### Metagenome-assembled genome analysis

The reads of each metagenome were mapped using minimap2 (Li 2018) with the -x and ava-ont parameters and then a *de novo* assembly was performed for each metagenome using the long reads assembler miniasm with default parameters (Li 2016).

#### BLAST

The assemblies of each metagenome were used as input to the command-line version of BLASTN (Camacho et al. 2009) against the bacterial tomato pathogens custom database described above and with the parameter of e-value set to less than or equal to 0.01. The top hit was determined to be the alignment with the longest length for each contig.

#### LINbase

The longest two contigs in each metagenome were used as input to LINbase at linbase.org (Tian et al. 2019) with the function “Identify using a genome sequence” to identify the pathogens at the strain level.

## Results

### Read-based pathogen identification after single-strain inoculation in the laboratory

Tomato plants inoculated with *Pto* isolate K40 (strain T1) in the laboratory showed bacterial speck symptoms four days after inoculation (Figure 1A), at which time DNA was extracted.

**Figure 1.**
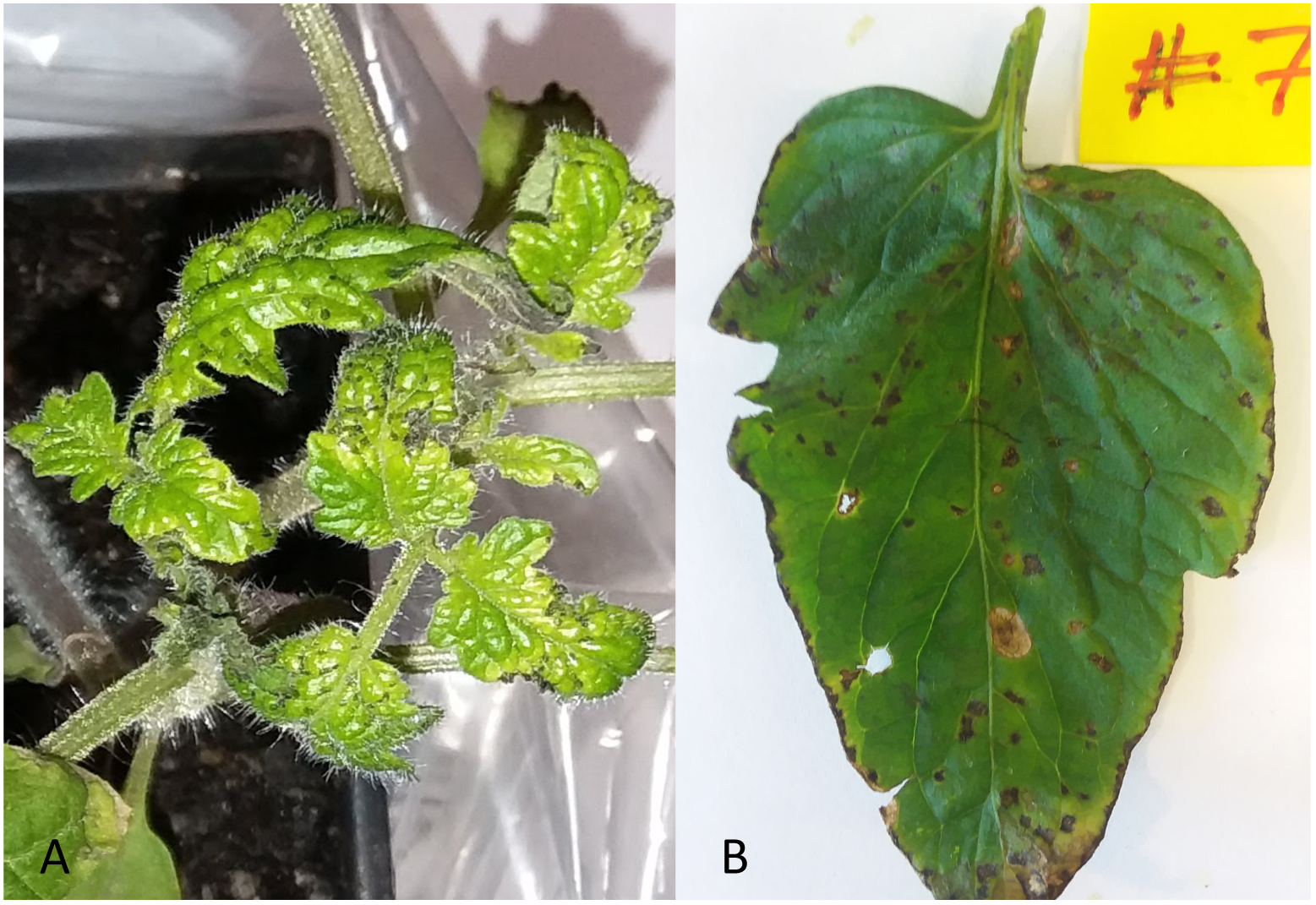
Diseased tomato plants (A) Symptoms caused by *Pseudomonas syringae* pv *tomato* isolate K40 (strain *Pto* T1) in a laboratory-inoculation assay and (B) Bacterial spot symptoms in naturally infected plants during a disease outbreak on the Eastern Shore of Virginia.

The quantity and quality of the extracted DNA is listed in Table 2. An entire MinION™ flow cell was used to sequence this sample (called L-K40). Of all the sequencing reads, 1,377,617 reads (approximately 60% of the total number of reads) were base-called after the run was completed using the guppy software. The base-called reads had a total length of approximately 4.2 Gigabases (Gbp) with the longest read measuring 66,000 bp (see more details about reads in Table 1).

**Table 2.**
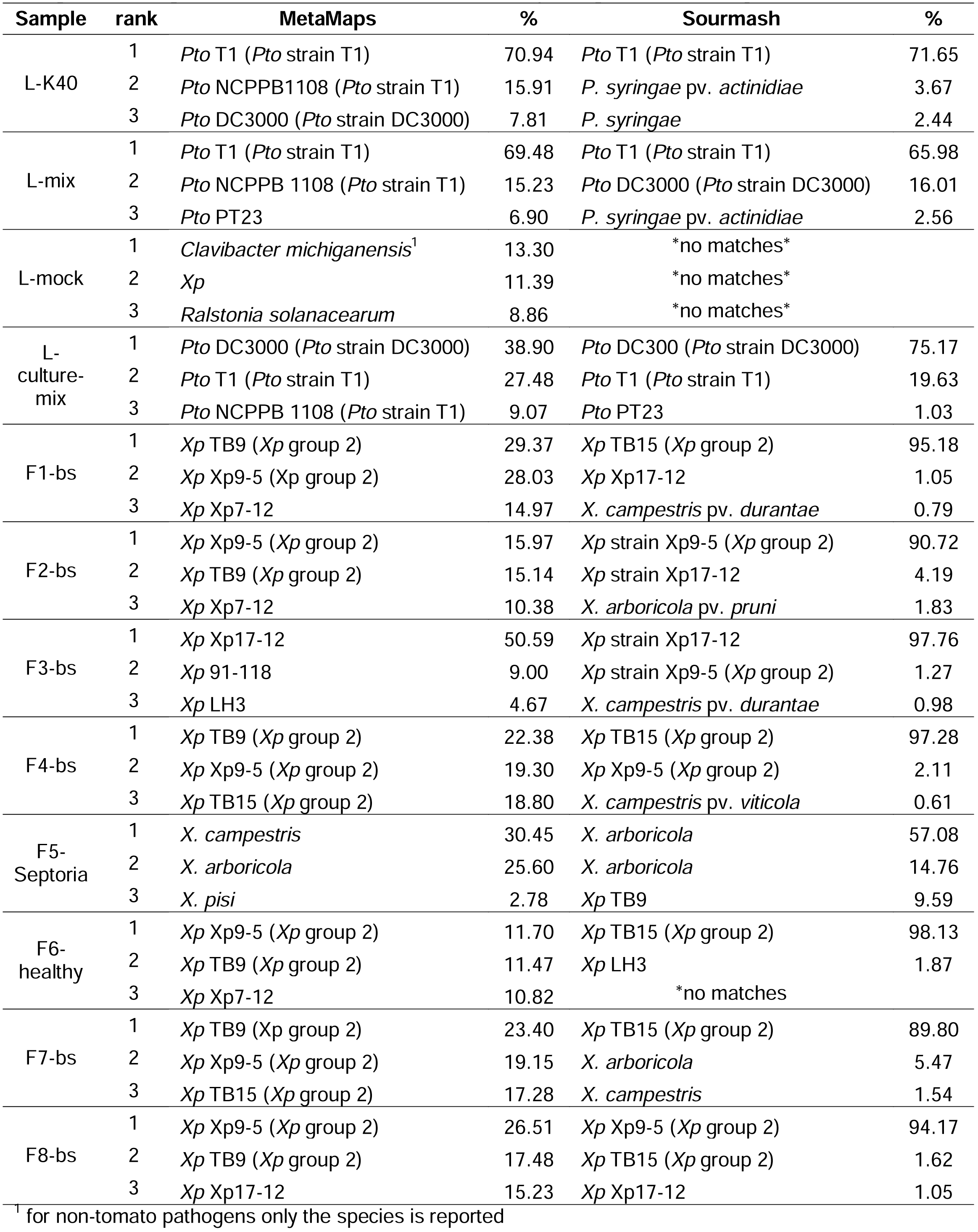
Relative abundance results (top three hits) obtained with MetaMaps and Sourmash using a custom genome database of bacterial tomato pathogens and closely related isolates.

The base-called reads were used as input to WIMP, which classified 89% of reads as of bacterial origin. Of these reads, WIMP identified 77.47% as *P. syringae* genomospecies 3, a genome similarity group of which *Pto* is a member. This genome similarity group was never validly published as a named species and is thus referred to with the number 3 instead of a name (Gardan et al. 1999). Also NCBI’s taxonomy database (Sayers et al. 2009) includes this taxon as *P. syringae* genomospecies 3. The next most abundant species were identified as *P. syringae* (9.39%), *P. cerasi* (2.09%), and *P. savastanoi* (1.60%). Figure 2 shows a screenshot of the WIMP result. The composition analysis is shown in Figure 3A (see Supplementary Table 2 for all relative abundance values for all composition analyses shown in Figure 3 and 4).

**Figure 2.**
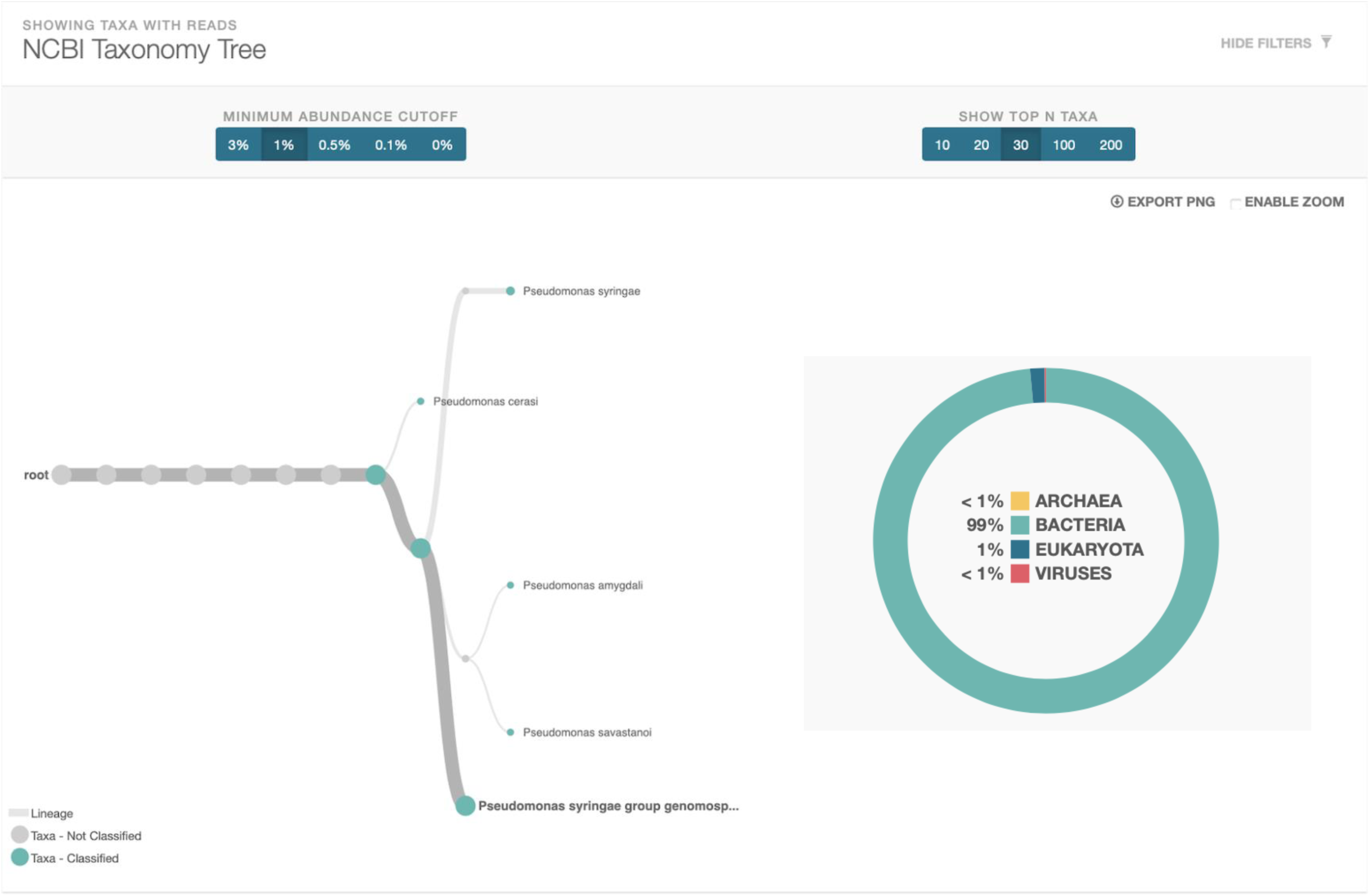
Screenshot of the WIMP taxonomy assignment for sample L-K40.

**Figure 3.**
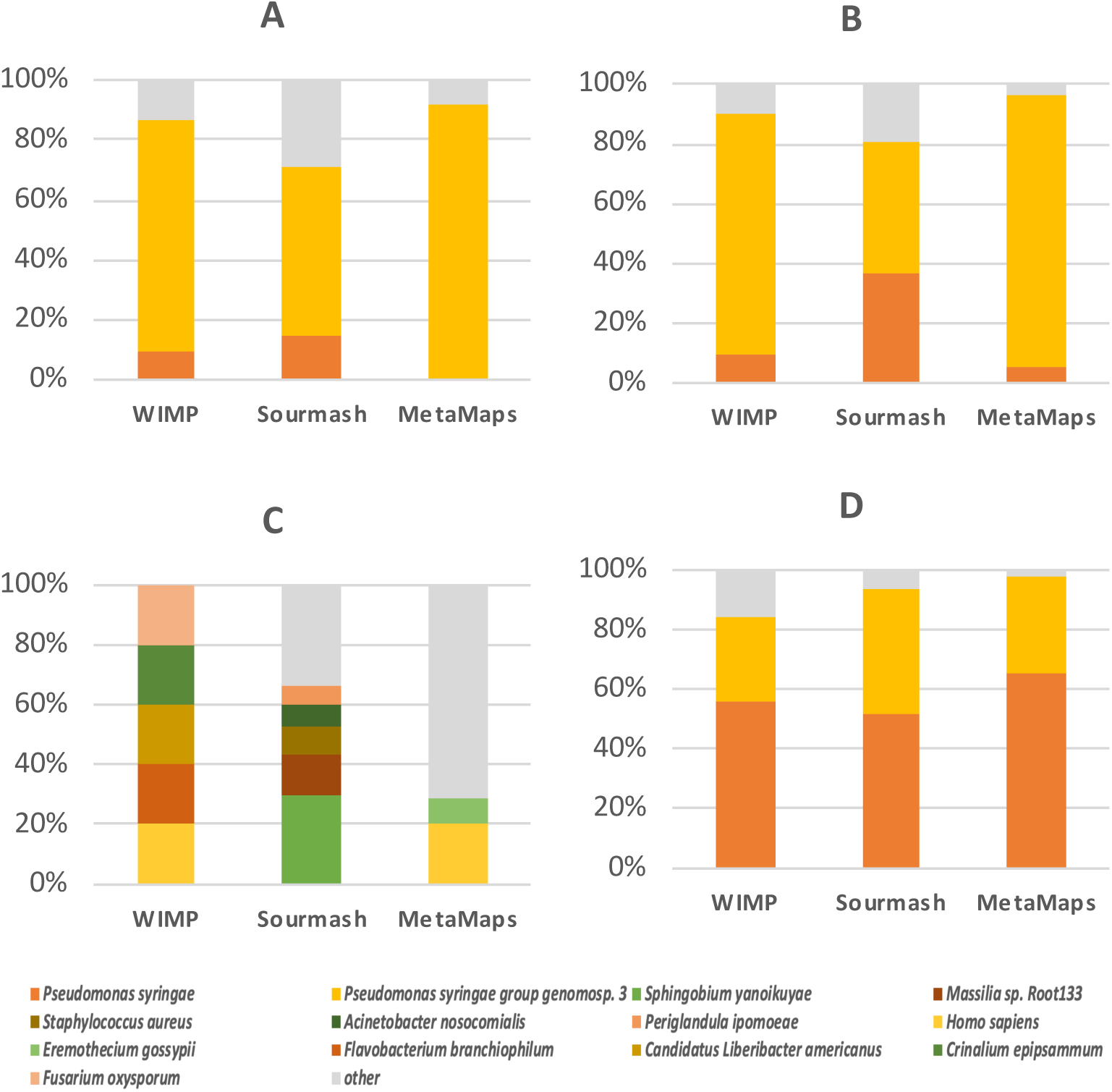
Bar graph showing the comparison of results at the species level using the read-based programs WIMP, Sourmash and MetaMaps. Each barplot corresponds to individual lab samples used in the study. A = L-K40, B = L-mix, C = L-mock, and D = L-culture-mix. Relative abundance values are expressed as percentages of all sequences classified as bacteria.

**Figure 4.**
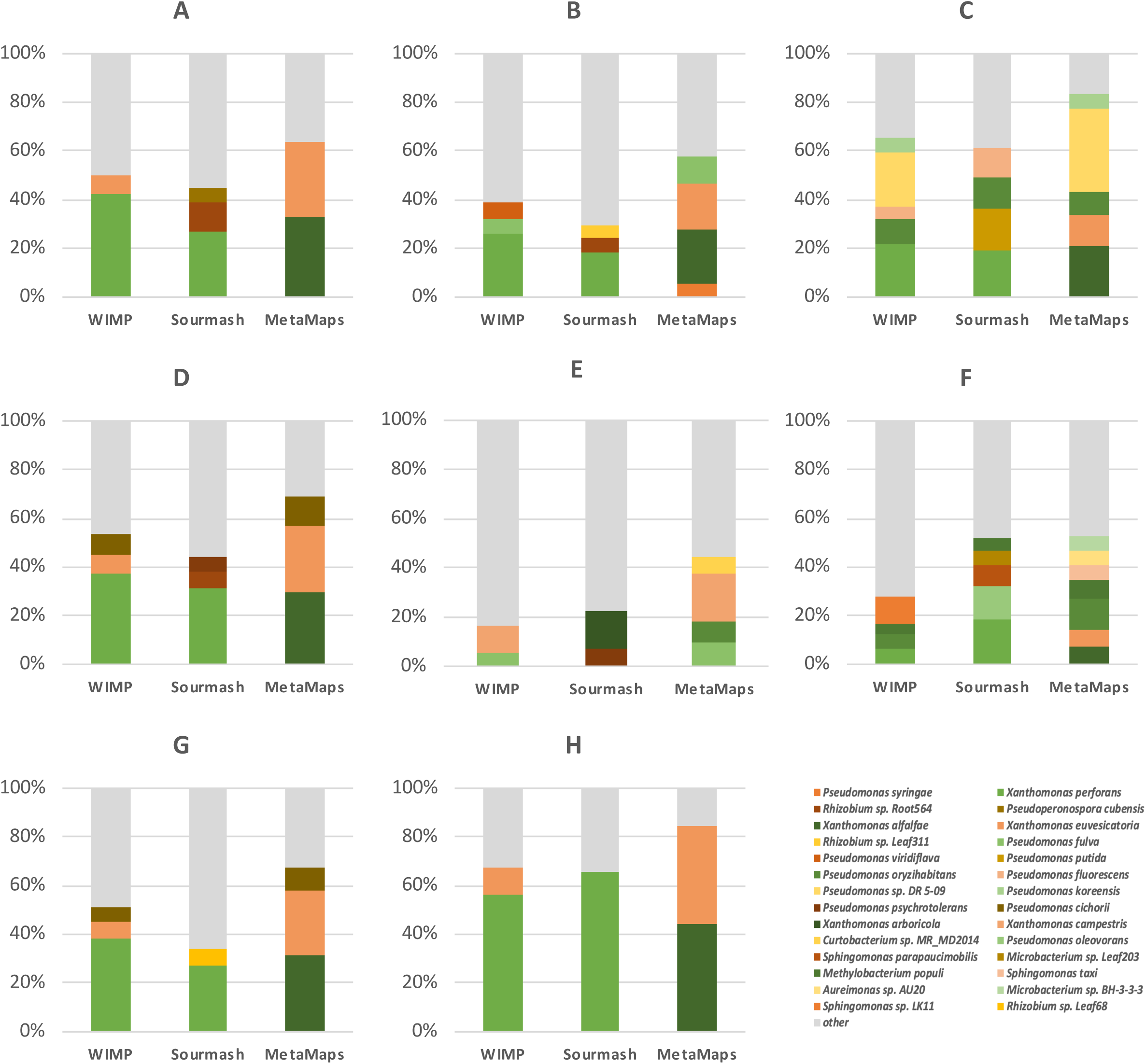
Bar graph showing the comparison of results at the species level using the read-based programs WIMP, Sourmash and MetaMaps. Each barplot corresponds to individual field samples used in the study. A = F1-bs, B = F2-bs, C = F3-bs, D = F4-bs, E = F5-Septoria, F = F6-healthy, G = F7-bs and H= F8-bs. Relative abundance values are expressed as percentages of all sequences classified as bacteria.

Next, the reads were used as input for composition analysis using Sourmash (Brown and Irber 2016) and MetaMaps (Dilthey et al. 2019) using the default reference libraries provided by these programs. Results are shown in Figure 3A. Sourmash identified 56.84% of the reads as *P. syringae* genomospecies 3 while MetaMaps identified over 91.53% of the reads as *P. syringae* genomospecies 3. Similarly to WIMP, both programs identified *P. syringae* as the next most abundant species (14.41% and 4.17%, respectively). All other species were found at a relative abundance of 2% or below. Therefore, WIMP, MetaMaps, and Sourmash all correctly identified the pathogen used in the inoculation as a member of *P. syringae* genomospecies 3. Supplementary Table 3 reports the run times for the three tools for this sample.

In an attempt to reach strain level resolution (not that WIMP is limited to species-level identification), we built Sourmash and MetaMaps custom reference libraries consisting of genome sequences of representative bacterial tomato pathogen isolates and closely related isolates that do not cause disease on tomato. The libraries included multiple isolates of the *Pto* strains DC3000 and T1 (Supplementary Table 2). When using these custom libraries, Sourmash identified 71.64% of the sequences in the sample as *Pto* isolate T1 (the isolate after which strain T1 is named) and the remaining sequences as other *P. syringae* isolates that are not pathogens of tomato (Table 2). Only 0.9% of the sequences were misidentified as *Pto* DC3000. MetaMaps in combination with the same custom library identified 70.93% as *Pto* isolate T1, 15.90% as *Pto* isolate NCPPB1108 (another isolate belonging to strain T1), and 7.81% as *Pto* isolate DC3000. Therefore, both Sourmash and MetaMaps identified most of the reads correctly as an isolate belonging to *Pto* strain T1 but Metamaps misidentified many more reads as *Pto* strain DC300 compared to Sourmash.

### Read-based pathogen identification after multi-strain inoculation in the laboratory

Next, we wanted to test the bioinformatics pipelines established with the single-strain inoculation by using a mixed inoculum consisting of the *Pto* isolate K40 (strain T1) and the *Pto* isolate DC3000 (strain DC3000) of *P. syringae* genomospecies 3 together with two additional isolates of the species *P. syringae* that do not cause disease on tomato: the bean pathogenic isolate *Psy* B728a and the non-pathogenic isolate *Psy* 642. DNA was again extracted on day four after inoculation and sequenced on an entire flow cell. All details for this sample (called L-mix) are listed in Table 1. Approximately 1 million reads of a total length of 4.2 Gbp were obtained with the longest read measuring 67,000 bp. Since this time 100% of reads were base-called, the number of base-called reads and the total length of reads were very similar to the single strain inoculation sample.

The caveat with this sample is that we did not know the relative abundance of the 4 isolates in the sample. However, since *Pto* isolates T1 and DC3000 are tomato pathogens while *Psy* isolates B728a and 642 are not, we expected that most sequences would be identified again as *P. syringae* genomospecies 3. In fact, WIMP identified 79.61% of all bacterial sequences (which constituted 95% of all reads) as *P. syringae* genomospecies 3 (Figure 3B), similar to the 77.47% identified in the single-strain inoculation sample. Compared to WIMP, Sourmash and MetaMaps showed the same trend as with the single strain inoculation sample: Sourmash found a lower relative abundance of *P. syringae* genomospecies 3 (43.24%) compared to WIMP and MetaMaps found a higher relative abundance compared to WIMP (91.09%) (Figure 3B).

Since both *Psy* isolates used in the inoculation belong to the species *P. syringae*, we expected a slightly higher relative abundance of *P. syringae* compared to the single strain inoculation sample. Interestingly, this expectation came true for Sourmash (36.87% versus 14.4%) but for WIMP and MetaMaps the relative abundance of *P. syringae* only increased marginally from 9.38% to 10.01% and from 4.17% to 5.39%, respectively (Figure 3B).

We then used the custom reference libraries of representative tomato pathogens to see if Sourmash and MetaMaps could distinguish isolate K40 (of strain T1) from isolate DC3000 (of strain DC3000). Sourmash did identify isolate T1 of strain T1 at a relative abundance of 65.98% and isolate DC3000 of strain DC3000 at a relative abundance of 16.01% (Table 2) while MetaMaps identified 84.71% of the reads as isolates that belong to strain T1 and 5.61% as isolate DC3000 (not shown in Table 2 since only the top three hits are shown for each sample).

Since we did not know the correct relative abundances of strains in this inoculated plant sample and could thus not determine how accurate the results were, we decided to sequence an additional sample (called L-culture-mix) that consisted of DNA extracted from an equal mixture of the same four strains after they were grown separately overnight in liquid culture. Approximately 54,000 reads of a total length of 150 Mbp were obtained on 1/6th of a flow cell with the longest read measuring 76,000 bp. WIMP classified 95% of the reads as bacterial. WIMP, MetaMaps, and Sourmash identified both, *P. syringae* and *P. syringae* genomospecies 3 in this sample, which we expected to be present at 50% each. WIMP over-estimated *P. syringae* compared to *P. syringae* genomospecies 3 (56% compared to 28%) and identified some other species at low relative abundance (Figure 3C). Metamaps also overestimated *P. syringae* compared to *P. syringae* genomospecies 3: 65.58% vs 32.19%. Sourmash came the closest to the expected 1 to 1 ratio finding 52.20% of *P. syringae* and 41.68% of *P. syringae* genomospecies 3 (Figure 3C). When using the custom reference libraries of tomato pathogens with MetaMaps and Sourmash, MetaMaps outperformed Sourmash since it identified DC3000 and T1 close to the expected 25% abundance: 38.89% and 27.48%, respectively (Table 2). Sourmash instead assigned a much higher abundance to strain DC3000 (75.1%) compared to strain T1 (19.63%) (Table 2).

Finally, we sequenced a tomato plant grown in the lab that was not inoculated with any pathogen (called sample L-mock). Since the DNA concentration of this sample was very low, only approximately 82,000 base-called reads were obtained on 1/7th of a flow cell with a total length of 103 Mb. The longest read was only 19,000 bp long. Only 8% of the reads were classified as bacterial showing that this lab-grown plant was not colonized by many bacteria, which was probably also the reason for the low DNA concentration. WIMP, Sourmash, and Metamaps provided very different results for this sample (Figure 3D). Importantly, as expected from a non-inoculated plant, none of the reads were identified by either of the three tools as *P. syringae* or *P. syringae* genomospecies 3.

### Read-based pathogen identification in naturally infected tomato field samples

After obtaining promising results in regard to strain-level identification with laboratory samples, we used DNA extracted from tomato field samples that were collected on the Eastern Shore of Virginia to test our pipelines with naturally infected plants (Table 1). The samples came from tomato plants that either showed symptoms of bacterial spot (samples F1-bs, F2-bs, F4-bs, F7-bs, F8-bs; see Figure 1B), symptoms of the fungal disease *Septoria* leaf spot (sample F5-Septoria) or no signs of any disease (F6-healthy). We also obtained one sample (F3-bs) with symptoms of bacterial spot but colonies that had been obtained from culturing bacteria from this plant had been found to be a mixture of colonies identified as either *Pseudomonas* or *Xanthomonas*.

DNA from all tomato field samples were barcoded and sequenced together with other samples by multiplexing them on the same flow cell. Therefore, the number of reads (between 35,923 for samples F6-healthy and 137,497 for F1-bs) and total read length (between 66 megabases (Mb) for F6-healthy and 588 Mb for F1-bs) for these samples were much lower compared to the laboratory samples (Table 1).

Detailed results for all samples are reported in Figure 4. Similarly to the lab-inoculated samples, the majority of reads in the field samples that had symptoms of bacterial disease were classified as bacteria by WIMP (between 78 and 81%). Importantly, WIMP and Sourmash agreed that *X. perforans* was the species with the highest relative abundance in these samples (between 25.82% and 56.44% for WIMP and between 18.51 and 66.01% for Sourmash) suggesting that *X. perforans* was the causative agent. Sample F3-bs, which had a mixed *Xanthomonas*/*Pseudomonas* infection based on culturing, was found by both WIMP and Sourmash to still be dominated by *X. perforans* (21.98% and 19.55% respectively) followed by either *P. oryzihabitans* (10.11%) and *P. fluorescens* (5.09%) based on WIMP or *P. putida* (16.98%) based on Sourmash. Therefore, the presence of a mixed infection was confirmed by both tools.

In contrast to the results from WIMP and Sourmash, MetaMaps identified *X. euvesicatoria* and *X. alfalfae* instead of *X. perforans* as the two species with the highest relative abundance in all samples with bacterial spot symptoms. This is because *X. perforans* was missing from the MetaMaps reference library.

Interestingly, even the non-symptomatic tomato sample (F6-healthy) was found to include *X. perforans* as the species with the highest relative abundance based on WIMP and Sourmash. However, the relative abundance values were lower (6.89% and 18.54%, respectively). This suggests that this plant might have been infected with *X. perforans* but was asymptomatic because of lower bacterial titer. This non-symptomatic sample also included a number of species at relatively high abundance that were rarely found in the samples with bacterial spot symptoms, for example, *P. oleovorans, Sphingomonas parapaucimobilis, Microbacterium sp.* Leaf203, and *Methylobacterium populi*.

The sample with *Septoria* leaf spot symptoms (F5-Septoria), probably infected by the plant pathogenic fungus *Septoria lycopersici*, carried a diverse bacterial population consisting of species in the genera *Pseudomonas, Xantomonas, Pantoea, Curtobacterium, Methylobacterium*, and *Sphingomonas*. No species in the fungal genus *Septoria* was included in any of the reference libraries and was thus not identified by any of the programs.

When we switched to Sourmash and MetaMaps using our custom database of representative bacterial tomato pathogens as reference libraries, *X. perforans* isolates TB9, TB15, and Xp9-5 were identified as the top hits in all plants with bacterial spot symptoms with the exception of F3-bs, which had the mixed *Pseudomonas*/*Xanthomonas* infection. In this sample, isolate Xp17-12 was identified by both Sourmash and MetaMaps as top hit. Interestingly, isolates TB9, TB15, and Xp9-5 are all members of the same intraspecific group, *X. perforans* group 2, based on core genome phylogeny (Schwartz et al. 2015), suggesting that the *X. perforans* strain infecting the tomatoes with bacterial spot symptoms on the Eastern Shore of Virginia was also a member of *X. perforans* group 2.

For sample F8-bs, we also isolated *Xanthomonas* bacteria to compare the results from the culture-independent read-based metagenomic approach with a culture-dependent genomic approach. DNA was extracted from two colonies and sequenced using Illumina HiSeq. The two genome sequences were assembled into 87 and 86 contigs, respectively, with a total length of 5,340,265 bp and 5,339,287 bp. We used the LINbase Web service for genome-based microbial identification and found isolate GEV1063 to be the best match for both genomes with 99.98% ANI and both genomes were identified by LINbase as members of *X. perforans* group 2, which is circumscribed in LINbase as an intraspecific taxon. Therefore, the culture-dependent genome-based identification confirmed the culture-independent read-based strain-level identification of *X. perforans* group 2 as the causative agent in sample F8-bs.

### Metagenome assembly-based pathogen identification

In parallel to the read-based pipelines described above, we also assembled each metagenomic sample using all reads that had a minimum length of 1,000 bp and that were identified by WIMP as bacterial. The results are summarized in Table 3. The non-inoculated tomato sample from the lab (L-mock), the healthy tomato sample from the field (F6-healthy), and the sample of the tomato plant with Septoria leaf spot (F5-Septoria) had the lowest number of contigs (between 4 and 9) with the shortest total length of contigs (between 21,390 bp and 122,956 bp). This was probably a result of the low number of bacterial reads in these samples (Table 1).

**Table 3.** Description of metagenomic assemblies.

| **Sample name** | **Total number of contigs** | **Total assembly length in bp** | **Mean contig length in bp** | **Longest contig in bp** | **2nd longest contig in bp** |
| --- | --- | --- | --- | --- | --- |
| L-K40 | 24 | 6,619,207 | 275,800 | 6,081,137 | 139,929 |
| L-mix | 73 | 8,669,208 | 118,756 | 6,126,095 | 118,770 |
| L-mock | 8 | 117,647 | 14,705 | 63,177 | 12,037 |
| L-culture-mix | 20 | 5,827,276 | 291,363 | 764,727 | 622,920 |
| F1-bs | 92 | 12,529,321 | 136,188 | 4,974,348 | 881,066 |
| F2-bs | 131 | 8,513,800 | 64,990 | 4,345,732 | 276,399 |
| F3-bs | 49 | 11,872,268 | 242,291 | 2,275,239 | 1,170,971 |
| F4-bs | 18 | 5,216,728 | 289,818 | 1,172,667 | 925,913 |
| F5-Septoria | 9 | 122,956 | 13,661 | 37,948 | 25,805 |
| F6-healthy | 4 | 21,390 | 5,347 | 8,488 | 7,900 |
| F7-bs | 35 | 5,666,575 | 161,902 | 5,038,472 | 56,441 |
| F8-bs | 10 | 5,319,638 | 531,963 | 2,680,062 | 2,212,039 |

The samples with symptoms of either bacterial speck or bacterial spot had a wide range in contig number and in the total length of contigs ranging from 10 to 131 contigs of a total length from 5.2 to 12.5Mbp. For our goal of identifying the causative agent in each symptomatic plant to strain level, we focused on the longest contigs in each sample since these contigs were the most likely to be of the causative pathogenic agents. It was very promising to see that in some of the symptomatic samples the longest contig was of a size similar to an entire bacterial genome, for example, 6.08Mbp in the tomato lab sample inoculated with Pto isolate K40 (L-K40), and 5.03Mbp for the field sample F7-bs showing bacterial spot symptoms (Table 3). We then used the genome alignment tool MUMmer (Marçais et al. 2018) to determine how much of the published genome sequences these contigs covered. We found that in the case of sample L-K40, the longest contig aligned with 93.92% of the published genome sequence of isolate K40. For F7-bis, the longest contig aligned with 95.52% of the published *X. perforans* genome of Xp8-16.

To obtain a preliminary identification of all contigs we used BLASTN (Camacho et al. 2009) in combination with our custom tomato pathogen database. The results were mostly in agreement with the reads-based analysis at the species level (Figure 5) but *X. euvesicatoria* was identified as species instead of *X. perforans* in some of the samples with bacterial spot.

**Figure 5.**
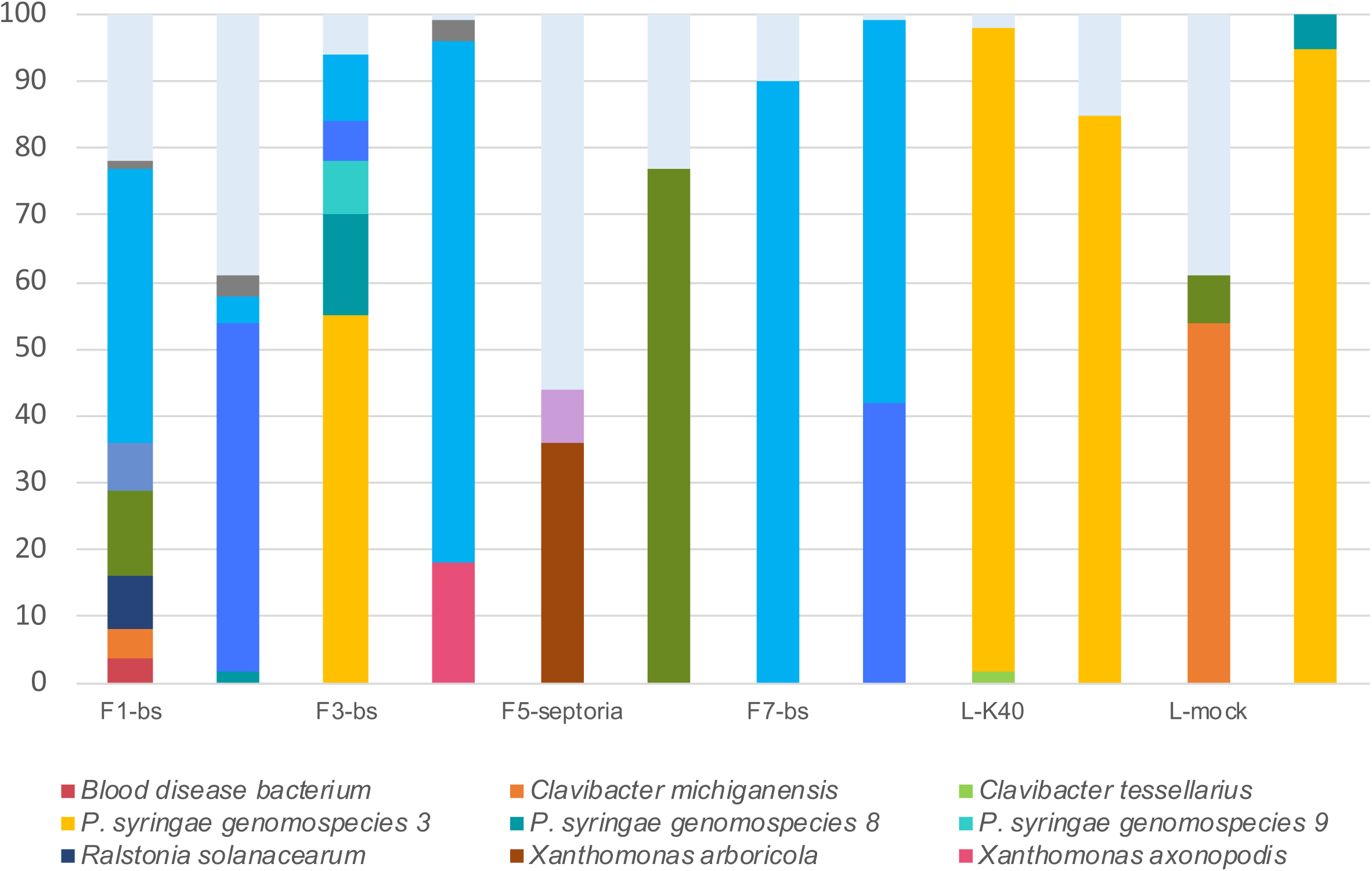
Relative genome percentage abundance for each sample based on BLASTN using contigs as query against a custom genome database. All hits were filtered to e-values less than or equal to 0.01 and the longest hit for each contig was considered to be the best hit.

To attempt identification of the longest contigs to strain level, we used these contigs as queries with the “Identify using a genome sequence” function in the LINbase Web service (Tian et al. 2019). Table 4 lists the results that were obtained for the longest two contigs (separately and merged) for each sample. When using the longest contig of the tomato plant inoculated with isolate K40 (of *Pto* strain T1), the *Pto* strain T1 isolate BAV1020 was the best hit but only with an ANI of 92.76% compared to the query sequence. However, based on a direct genome sequence comparison, the two genomes are over 99.75% identical to each other. Since we know that isolate K40 was used as inoculum, the discrepancy between the two ANI value is necessarily a result of the high error rate of the MinION™ sequencer.

**Table 4.** LINbase results for two longest contigs

| Sample | Longest contig (ANI %) | Taxon membership of longest contig | Second longest contig (ANI %) | Taxon membership of second longest | Two longest contigs merged (ANI %) | Taxon membership of merged contigs |
| --- | --- | --- | --- | --- | --- | --- |
| L-K40 | *Pto* BAV1020 (92.766) | *Pto* strain T1 | NA | NA | *Pto* BAV1020 (92.761) | *Pto* strain T1 |
| L-mix | *Pto* BAV1020 (92.731) | *Pto* strain T1 | NA | NA | *Pto* NYS-T1 (92.769) | *Pto* strain T1 |
| L-culture-mix | *Ps* 642 (93.368) | *Ps* | *Ps* UB0390 (93.408) | *Ps* | *Pc* ICMP19117 (93.315) | *Pseudomonas* |
| F1-bs | *Xp* Xp10-13 (94.625) | *Xp* group 2 | NA | NA | *Xp* GEV1063 (94.613) | *Xp* group 2 |
| F2-bs | *Xp* GEV2117 (94.236) | *Xp* group 2 | *Xp* 91-118 (94.478) | *Xp* | *Xp* GEV2117 (94.255) | *Xp* group 2 |
| F3-bs | *Pf* Pf0-1 (89.669) | *Pseudomonas* | *Pf* Pf0-1 (89.710) | *Pseudomonas* | *Pf* Pf0-1 (89.675) | *Pseudomonas* |
| F4-bs | *Xp* 91-118 (94.263) | *Xp* | *Xp* 91-118 (94.501) | *Xp* | *Xp* 91-118 (94.369) | *Xp* |
| F7-bs | *Xp* Xp8-16 (94.464) | *Xp* group 2 | NA | NA | *Xp* GEV2116 (94.360) | *Xp* group 2 |
| F8-bs | *Xp* Xp10-13 (93.322) | *Xp* group 2 | *Xp* GEV2117 (93.271) | *Xp* group 2 | *Xp* Xp10-13 (93.352) | *Xp* group 2 |
| BAV6163 | *Xp* GEV1063 (99.976) | *Xp* group 2 |  |  |  |  |
| BAV6164 | *Xp* GEV1063 (99.98) | *Xp* group 2 |  |  |  |  |

*Ps* = *Pseudomonas syringae* *Pf* = *Pseudomonas fluorescens* *Pc* = *Pseudomonas congelans* *Xp* = *Xanthomonas perforans*
NA – Not available, second contig too short for identification

For the tomato plant inoculated with the four-strain mix, the longest contig was again identified as *Pto* strain T1 based on the best hit to *Pto* isolate T1 with an ANI value of 92.73%. No contig of significant length was identified as *Pto* isolate DC3000. Since the genomes of *Pto* isolates DC3000 and T1 are over 98.5% identical to each other, the longest contig of this sample was probably assembled from a combination of DC3000 and T1 reads, which could not be distinguished from each other also because of the high error rate of the MinION™ sequencer.

For the longest contigs in the tomato field samples that showed bacterial spot symptoms, different isolates of *X. perforans* were the best hits: Xp8-16, Xp10-13, GEV1063, and GEV2116 (Table 4). These isolates belong to *X. perforans* group 2 (Schwartz et al. 2015) and are thus in line with the read-based results described above. Only the second-longest contig in sample F2-bs and the two longest contigs in sample F4-bs contradicted the read-based results: *X. perforans* isolate 91-118, a member of *X. perforans* group 1B (Schwartz et al. 2015), was the best hit for these contigs.

Since for sample F8-bs we also had the genome sequences of the two cultured isolates (see previous section), we could again directly compare the metagenomic assembly-based approach with the culture-dependent genomic approach. Although there was no difference in the identification results themselves since the best matches in LINbase for both approaches were isolates of *X. perforans* group 2, the ANI between the longest contig of F8-bs and the most similar genome in LINbase was only 93.35% while the ANI between the genome sequences of the isolated colonies and their most similar genome in LINbase was 99.98%. As with the lab-inoculated sample L-K40, this difference in ANI was probably again due to the high error rate of the MinION™ and was the reason we could not directly identify the causative agent as a member of *X. perforans* group 2.

## Discussion

Sensitive detection and precise identification of pathogens in real time directly from symptomatic organisms, or even better from infected but still asymptomatic organisms, without the need for pathogen isolation and culturing, is the ultimate goal in control and prevention of infectious diseases of humans, animals, and plants.

As a step towards this goal in plant pathology, here we used the ONT MinION™ for precise identification of two bacterial tomato pathogens by sequencing metagenomic DNA directly extracted from symptomatic plants and analyzing the obtained sequences with a set of different tools and databases. However, we neither attempted to maximize sensitivity of detection nor to minimize the time necessary for identification.

Several other reports describing the use of the MinION™ in culture-independent metagenomic DNA sequencing for plant pathogen identification have recently been published. Most of these reports either focused on species-level identification (Hu et al. 2019) and/or on accelerating the identification protocol (Loit et al. 2019). Only one report focused on strain-level identification but after polymerase chain reaction with primers specific to loci of a single pathogen species, which increased the sensitivity of detection and resolution of identification but restricts the approach to a single pathogen species at the time (Radhakrishnan et al. 2019). Our goal instead was to develop an experimental and bioinformatics pipeline that can be used for any bacterial plant pathogen, and, with modifications, possibly for fungal and oomycete pathogens as well.

The first critical step in metagenomic-based pathogen identification is DNA extraction. There are mainly two possibilities: extracting DNA directly from plant tissue or extracting DNA from water used to wash the plant (after sonication to help dislocate the pathogen from the tissue). The first approach has the advantage that large quantities of high-quality DNA can be extracted. The obvious disadvantage is that a large fraction of the extracted DNA is plant DNA. The second approach is the approach we decided to use since it is widely used for plant microbiome analysis, for example (Ottesen et al. 2013). Based on the results from our DNA sequence analysis, this approach allowed us to obtain DNA that was over 80% of bacterial origin for the naturally infected tomato field samples and over 90% of bacterial origin for the artificially inoculated tomato plants grown in the laboratory. This value was as high as the fraction of bacterial DNA when extracting DNA directly from a bacterial culture. Therefore, we conclude that for metagenome-based identification of bacterial foliar pathogens in symptomatic plant tissue extracting DNA from wash water after sonication is an excellent solution. Importantly, even the wash water of our healthy field sample still contained 30% of bacterial DNA, making this approach possibly still a good choice even for asymptomatic leaves with relatively low bacterial titers.

Because in this project we were not interested in speed, we used the slower, higher yielding DNA sequencing library preparation protocol, as suggested by ONT, without significant modifications. Also for the sequencing protocol itself, we followed ONT’s instructions without modifications. The first critical step after sequencing the DNA, is base-calling, which is the process of translating the raw electrical signals measured by the MinION™ into nucleotide sequences. Since base-calling is computationally intensive and takes longer than sequencing itself, base-calling needed to be completed after the sequencing runs themselves were completed. We used the ONT Guppy base-calling tool without any polishing.

The actual assignment of sequencing reads to specific bacterial species and strains was done using a total of five tools: 1. ONT’s WIMP software with graphical user interface, which is intuitive to use and uses the software Centrifuge (Kim et al. 2016) to rapidly identify and assign taxonomy to the reads coming from the sequencing base calling in real-time, 2. the command-line tool Sourmash (Brown and Irber 2016) that computes hash sketches from DNA sequences and includes k-mer based taxonomic classification for genomic and metagenomic analysis, 3. the command line tool MetaMaps (Dilthey et al. 2019) which uses approximate mapping algorithm to map long-read metagenomic sequences to comprehensive databases, 4. the command line version of BLASTN (Camacho et al. 2009) was used to speed up the identification of pathogens after metagenome assembly with a custom-built database, 5. assembly of metagenomes obtained by minimap2 and miniasm (Li 2016) followed by taxonomy assignment of the two longest contigs obtained by LINbase (Tian et al. 2019). Moreover, Sourmash and MetaMaps were used both with default and custom libraries.

For species-level identification, the three read-based tools performed similarly well with the lab samples in regard to accuracy with Sourmash coming the closest to the expected 1: 1 ratio of *P. syringae* genomospecies 3: *P. syringae* in the sample L-culture-mix. For the field samples, the absence of *X. perforans* in the MetaMaps default reference library did not allow MetaMaps to identity *X. perforans* while WIMP and Sourmash performed similarly well. Both identified *X. perforans* as the most abundant species in all samples with bacterial spot symptoms.

As for run time, only WIMP is set up to provide real-time results starting minutes after runs are initiated and results are updated as more sequencing reads are base-called. However, since base-calling cannot keep up with the amount of raw data that is being generated during a run, WIMP needs to be re-run when base-calling is completed after a run ends in order to analyze all data. This took over 36 hours for our largest sample, L-K40 (Supplementary Table 3). The advantage is that users do not need any significant local computing resources to do this since WIMP runs on ONT’s cloud. For the same L-K40 sample, it took Sourmash only 35 minutes to calculate the k-mer signature and perform species-level classification while Metamaps completed the same run in 6-8 hours. Both tools were run on Virginia Tech’s ARC high-performance computing system. Therefore, Sourmash is significantly faster than MetaMaps and WIMP but still requires significant computing resources.

In regard to ease of use, WIMP cannot be beaten because of its intuitive graphical user interface. Although both Sourmash and Metamaps are command-line tools, Sourmash beats Metamaps because of the extensive tutorials provided on the Sourmash website. The added ease of making custom reference libraries and adding genomes to existing libraries also makes Sourmash more user-friendly compared to MetaMaps, which requires NCBI taxIDs (or creation of custom taxIDs) for all genomes in custom reference libraries.

Assembling reads into contigs before identification did not provide any advantages for species-level identification since species-level identification was successful with read-based tools and read-based identification is generally faster since it does not require prior assembly of reads into contigs. However, this advantage of speed may diminish with an increasing number of reads since mapping of a smaller number of assembled contigs might be faster than mapping a large number of reads individually.

For strain-level identification, WIMP cannot be used since it only reaches species-level resolution. When comparing MetaMaps with Sourmash, MetaMaps misidentified a larger number of reads as strain *Pto* DC3000 compared to Sourmash in the single strain inoculation sample L-K40, which we knew did not contain any DNA of strain *Pto* DC3000. Instead in the sample L-culture-mix with known equal concentrations, it was Sourmash that overestimated strain *Pto* DC3000 compared to strain *Pto* T1. For field sample F8-bs for which we had also a culture-dependent result indicating *X. perforans* group 2 as causative agent, both software identified the same best hit in the custom database that was also a member of *X. perforans* group 2. Therefore, we conclude that Sourmash and MetaMaps did equally well in regard to strain accuracy. In regard to run time, Sourmash’s run time increased to 1-3 hours when using a k-mer size of 51, which is required for strain-level identification. Run time for MetaMaps decreased to 3-4 hours because of the smaller size of the custom library in comparison to default databases. However, Sourmash still performed better than MetaMaps in regard to computation time.

The challenge when using either Sourmash or MetaMaps for strain-level identification is that we had to interpret the results based on prior knowledge of which isolates in our custom database belonged to which pathogen strain. For example, only by checking Figure 1 in (Schwartz et al. 2015), were we able to identify the best matches found by Sourmash and MetaMaps in our custom database as members of *X. perforans* group 2. Moreover, a best match with an isolate that belongs to a certain strain, or any other group or taxon for that matter, still does not necessarily mean that the query is a member of the same group as well. To make such a conclusion, it is necessary to determine (1) the genomic breadth of the group, for example, 99.75% for *X. perforans* group 2, and (2) the genomic distance of the query to a representative member of that group with this distance needing to be smaller than the genomic breadth of the group. Alternatively, a phylogenetic analysis could be performed to determine if the unknown is a member of the clade that corresponds to the specific group. Because species have a standard genomic breadth of 95% ANI, WIMP, Sourmash, and Metamaps can infer species membership from metagenomic reads relatively easily. However, strains (and any other group smaller than a species) do not have a standard ANI breadth. Therefore, Sourmash and MetaMaps would need to be given genomic circumscriptions of strains as part of the reference library information in order to precisely assign reads to strains.

Since the MinION™ outputs long reads, we were surprisingly successful in assembling reads into contigs almost as long as entire bacterial genomes, which could then be used for genome-based identification. We specifically developed the LINbase Web service for identifying microbes as members of taxa at any genomic breadth below the rank of genus (Tian et al. 2019) and we had circumscribed both *Pto* strain T1 and *X. perforans* group 2 as taxa in LINbase with genomic breadths of 99.75% and 99.9% ANI, respectively. Therefore, we should have been able to avoid the problem that we had with read-based identification. However, the challenge that arose with this approach was that because of the high error rate of the MinION™, the ANI between all query contigs and their best matches in LINbase were below 95%. This was true even for the longest contig in sample L-K40, which had been inoculated with strain *Pto* T1 isolate K40. Therefore, the longest contig in this sample should have had an almost 100% match in LINbase with the genome of isolate K40 and other isolates that belong to strain T1. However, the ANI between this contig and the best match in LINbase was only 92.76%. Therefore, using the metagenome-assembled contigs did not allow us to identify the pathogens as members of the strains circumscribed in LINbase because the MinION™ error rate lowered the ANI between the query contig and the best match to below the genomic breath of the circumscribed taxon. Being aware of the high error rate, we were still able to extrapolate from the best match in LINbase the identity of the correct strain. However, such a result can only be considered putative or preliminary.

In conclusion, using either the Sourmash and MetaMaps tools for read-based strain identification or LINbase for assembly-based strain-level identification, putative strain-level identification was possible and was confirmed by culture-dependent genome-based identification. However, it was impossible to reach high-confidence strain-level identification because of the absence of appropriate strain-level databases for the read-based tools and because of the high error rate of the MinION™when using assembly-based identification. Considering the large and active user community of the MinION™ sequencer and the continued development of new versions of the MinION™, we expect improvements in both, tool development for read-based identification, and improvements in the precision at which the MinION™ can distinguish nucleotides from each other and/or base-calling algorithms, which should ultimately lower the currently high error rate. At this point, we consider culture-independent metagenomic sequencing with the MinION™ an excellent approach to obtain results when high confidence strain-level identification is not required or when a culture-dependent genome-based identification is used as a follow-up.

## Author contributions

BAV and SL developed the project. MEML performed most of the wet-lab experiments. MAF and PS did most of the bioinformatics analyses. SY contributed to the wet-lab experiments. LT and CH, under supervision from BAV and LSH, developed LINbase. BAV, with contributions from MEML, MAF, PS, and SL wrote the manuscript. All authors read and approved the final version of the manuscript.

## Conflict of Interest

LINbase uses the trademarks Life Identification Number^®^ and LIN^®^, which are registered by This Genomic Life, Inc. LSH and BAV report in accordance with Virginia Tech policies and procedures and their ethical obligation as researchers that they have a financial interest in This Genomic Life, Inc. Therefore, their financial interests may be affected by the research reported in this manuscript. They have disclosed those interests fully to Virginia Tech, and they have in place an approved plan for managing any potential conflicts arising from this relationship.

## Funding

This study was supported by the College of Agriculture and Life Sciences at Virginia Polytechnic Institute and State University and by the National Science Foundation (IOS-1754721). Funding to BAV and SL was also provided in part by the Virginia Agricultural Experiment Station and the Hatch Program of the National Institute of Food and Agriculture, US Department of Agriculture.

## Acknowledgements

The authors acknowledge Advanced Research Computing (ARC) at Virginia Tech for providing computational resources and technical support that have contributed to the results reported within this paper. URL: http://www.arc.vt.edu

## Supplementary Tables

**Supplementary Table 1.** List of genomes used in the custom database.

**Supplementary Table 2.** Relative abundance values at the species level for all samples obtained with WIMP, Sourmash, and MetaMaps.

**Supplementary Table 3.** Example run times for WIMP, Sourmash, and MetaMaps.

## Literature cited

Almeida, N. F., Yan, S., Cai, R., Clarke, C. R., Morris, C. E., Schaad, N. W., et al. 2010. PAMDB, a multilocus sequence typing and analysis database and website for plant-associated microbes. Phytopathology. 100:208–215.

Andrews, S. 2010. Babraham bioinformatics-FastQC a quality control tool for high throughput sequence data. URL: https://www.bioinformatics.babraham.ac.uk/projects/fastqc/(accessed 06. 12. 2018). Available at: https://www.bioinformatics.babraham.ac.uk/projects/fastqc/.

Badial, A. B., Sherman, D., Stone, A., Gopakumar, A., Wilson, V., Schneider, W., et al. 2018. Nanopore Sequencing as a Surveillance Tool for Plant Pathogens in Plant and Insect Tissues. Plant Disease. 102:1648–1652 Available at: http://dx.doi.org/10.1094/pdis-04-17-0488-re.

Brown, C. T., and Irber, L. 2016. sourmash: a library for MinHash sketching of DNA. J. Open Source Software. 1:27.

Bushnell, B. 2015. BBMap. Available at: https://sourceforge.net/projects/bbmap/.

Cai, R., Lewis, J., Yan, S., Liu, H., Clarke, C. R., Campanile, F., et al. 2011. The plant pathogen *Pseudomonas syringae* pv. *tomato* is genetically monomorphic and under strong selection to evade tomato immunity. PLoS Pathog. 7:e1002130.

Camacho, C., Coulouris, G., Avagyan, V., Ma, N., Papadopoulos, J., Bealer, K., et al. 2009. BLAST+: architecture and applications. BMC Bioinformatics. 10:421.

Chalupowicz, L., Dombrovsky, A., Gaba, V., Luria, N., Reuven, M., Beerman, A., et al. 2019. Diagnosis of plant diseases using the Nanopore sequencing platform. Plant Pathol. 68:229–238.

Clarke, C. R., Cai, R., Studholme, D. J., Guttman, D. S., and Vinatzer, B. A. 2010. *Pseudomonas syringae* strains naturally lacking the classical *P. syringae hrp/hrc* Locus are common leaf colonizers equipped with an atypical type III secretion system. Mol. Plant. Microbe. Interact. 23:198–210.

Dijkshoorn, L., Ursing, B. M., and Ursing, J. B. 2000. Strain, clone and species: comments on three basic concepts of bacteriology. J. Med. Microbiol. 49:397–401.

Dilthey, A. T., Jain, C., Koren, S., and Phillippy, A. M. 2019. Strain-level metagenomic assignment and compositional estimation for long reads with MetaMaps. Nat. Commun. 10:3066.

Fang, Y., and Ramasamy, R. P. 2015. Current and Prospective Methods for Plant Disease Detection. Biosensors. 5:537–561.

Feil, H., Feil, W. S., Chain, P., Larimer, F., DiBartolo, G., Copeland, A., et al. 2005. Comparison of the complete genome sequences of *Pseudomonas syringae* pv. syringae B728a and pv. tomato DC3000. Proc. Natl. Acad. Sci. U. S. A. 102:11064–11069.

Gardan, L., Shafik, H., Belouin, S., Broch, R., Grimont, F., and Grimont, P. A. 1999. DNA relatedness among the pathovars of *Pseudomonas syringa*e and description of *Pseudomonas tremae* sp. nov. and *Pseudomonas cannabina* sp. nov. (ex Sutic and Dowson 1959). Int. J. Syst. Bacteriol. 49 Pt 2:469–478.

Hu, Y., Green, G. S., Milgate, A. W., Stone, E. A., Rathjen, J. P., and Schwessinger, B. 2019. Pathogen Detection and Microbiome Analysis of Infected Wheat Using a Portable DNA Sequencer. Phytobiomes Journal. 3:92–101.

Jain, M., Olsen, H. E., Paten, B., and Akeson, M. 2016. The Oxford Nanopore MinION: delivery of nanopore sequencing to the genomics community. Genome Biol. 17:239.

Jones, J. B., Lacy, G. H., Bouzar, H., Stall, R. E., and Schaad, N. W. 2004. Reclassification of the Xanthomonads associated with bacterial spot disease of tomato and pepper. Syst. Appl. Microbiol. 27:755–762.

Juul, S., Izquierdo, F., Hurst, A., Dai, X., Wright, A., Kulesha, E., et al. 2015. What’s in my pot? Real-time species identification on the MinION. bioRxiv.: 030742.

Kim, D., Song, L., Breitwieser, F. P., and Salzberg, S. L. 2016. Centrifuge: rapid and sensitive classification of metagenomic sequences. Genome Res. 26:1721–1729.

Konstantinidis, K. T., and Tiedje, J. M. 2005. Genomic insights that advance the species definition for prokaryotes. Proc. Natl. Acad. Sci. U. S. A. 102:2567–2572.

Li, H. 2018. Minimap2: pairwise alignment for nucleotide sequences. Bioinformatics. 34:3094–3100.

Li, H. 2016. Minimap and miniasm: fast mapping and de novo assembly for noisy long sequences. Bioinformatics. 32:2103–2110.

Loit, K., Adamson, K., Bahram, M., Puusepp, R., Anslan, S., Kiiker, R., et al. 2019. Relative performance of Oxford Nanopore MinION vs. Pacific Biosciences Sequel third-generation sequencing platforms in identification of agricultural and forest pathogens. bioRxiv.: 592972 Available at: https://www.biorxiv.org/content/10.1101/592972v1.abstract [Accessed September 8, 2019].

Marçais, G., Delcher, A. L., Phillippy, A. M., Coston, R., Salzberg, S. L., and Zimin, A. 2018. MUMmer4: A fast and versatile genome alignment system. PLoS Comput. Biol. 14:e1005944.

Mechan-Llontop, M. E., Tian, L., Bernal-Galeano, V., Reeves, E., Hansen, M. A., Bush, E., et al. 2019. Assessing the potential of culture-independent 16S rRNA microbiome analysis in disease diagnostics: the example of *Dianthus gratianopolitanus* and *Robbsia andropogonis*. European Journal of Plant Pathology. Available at: http://dx.doi.org/10.1007/s10658-019-01850-8 [Accessed September 16, 2019].

MinION brochure. 2019a. Oxford Nanopore Technologies. Available at: http://nanoporetech.com/resource-centre/minion-brochure [Accessed September 14, 2019].

MinION brochure. 2019b. Oxford Nanopore Technologies. Available at: http://nanoporetech.com/resource-centre/minion-brochure [Accessed September 14, 2019].

Nadon, C., Van Walle, I., Gerner-Smidt, P., Campos, J., Chinen, I., Concepcion-Acevedo, J., et al. 2017. PulseNet International: Vision for the implementation of whole genome sequencing (WGS) for global food-borne disease surveillance. Euro Surveill. 22 Available at: http://dx.doi.org/10.2807/1560-7917.ES.2017.22.23.30544.

Ottesen, A. R., González Peña, A., White, J. R., Pettengill, J. B., Li, C., Allard, S., et al. 2013. Baseline survey of the anatomical microbial ecology of an important food plant: *Solanum lycopersicum* (tomato). BMC Microbiol. 13:114.

Radhakrishnan, G. V., Cook, N. M., Bueno-Sancho, V., Lewis, C. M., Persoons, A., Mitiku, A. D., et al. 2019. MARPLE, a point-of-care, strain-level disease diagnostics and surveillance tool for complex fungal pathogens. BMC Biology. 17 Available at: http://dx.doi.org/10.1186/s12915-019-0684-y.

Rees-George, J., Vanneste, J. L., Cornish, D. A., Pushparajah, I. P. S., Yu, J., Templeton, M. D., et al. 2010. Detection of *Pseudomonas syringae* pv. actinidiae using polymerase chain reaction (PCR) primers based on the 16S-23S rDNA intertranscribed spacer region and comparison with PCR primers based on other gene regions. Plant Pathology. 59:453–464 Available at: http://dx.doi.org/10.1111/j.1365-3059.2010.02259.x.

Sayers, E. W., Barrett, T., Benson, D. A., Bryant, S. H., Canese, K., Chetvernin, V., et al. 2009. Database resources of the National Center for Biotechnology Information. Nucleic Acids Res. 37:D5–15.

Schwartz, A. R., Potnis, N., Timilsina, S., Wilson, M., PatanÃ©, J., Martins, J., et al. 2015. Phylogenomics of *Xanthomonas* field strains infecting pepper and tomato reveals diversity in effector repertoires and identifies determinants of host specificity. Frontiers in Microbiology. 6 Available at: http://dx.doi.org/10.3389/fmicb.2015.00535.

Tedersoo, L., Drenkhan, R., Anslan, S., Morales-Rodriguez, C., and Cleary, M. 2019. High-throughput identification and diagnostics of pathogens and pests: Overview and practical recommendations. Molecular Ecology Resources. 19:47–76 Available at: http://dx.doi.org/10.1111/1755-0998.12959.

Tian, L., Huang, C., Heath, L. S., and Vinatzer, B. A. 2019. LINbase: A Web service for genome-based identification of microbes as members of crowdsourced taxa. bioRxiv. Available at: https://www.biorxiv.org/content/10.1101/752212v1.abstract.

Tinivella, F., Gullino, M. L., and Stack, J. P. 2008. The Need for Diagnostic Tools and Infrastructure. In Crop Biosecurity, Springer Netherlands, p. 63–71.

Wick, R. R., Judd, L. M., Gorrie, C. L., and Holt, K. E. 2017. Unicycler: Resolving bacterial genome assemblies from short and long sequencing reads. PLoS Comput. Biol. 13:e1005595.

Williamson, L., Nakaho, K., Hudelson, B., and Allen, C. 2002. *Ralstonia solanacearum* Race 3, Biovar 2 Strains Isolated from Geranium Are Pathogenic on Potato. Plant Dis. 86:987–991.

Yan, S., Liu, H., Mohr, T. J., Jenrette, J., Chiodini, R., Zaccardelli, M., et al. 2008. Role of Recombination in the Evolution of the Model Plant Pathogen *Pseudomonas syringae* pv. *tomato* DC3000, a Very Atypical Tomato Strain. Applied and Environmental Microbiology. 74:3171–3181 Available at: http://dx.doi.org/10.1128/aem.00180-08.

